# MYH10 governs adipocyte function and adipogenesis through its interaction with GLUT4

**DOI:** 10.1101/2021.12.07.471535

**Authors:** N Kislev, L Mor-Yossef Moldovan, R Barak, M Egozi, D Benayahu

**Affiliations:** Department of Cell and Developmental Biology, Sackler School of Medicine, Tel Aviv University, Tel Aviv 6997801, Israel

**Author notes:** Correspondence Prof. Dafna Benayahu, Department of Cell and Developmental Biology, Sackler School of Medicine, Tel Aviv University, Tel Aviv 6997801, Israel. Equal contribution (N Kislev, L Mor-Yossef Moldovan).

**Keywords:** MYH10, GLUT4, PKCζ, adipogenesis, cytoskeleton organization, insulin signaling

## Abstract

Adipocyte differentiation is dependent on cytoskeletal remodeling processes that determine and maintain cellular shape and function. In turn, cytoskeletal proteins contribute to the filament-based network responsible for controlling adipocyte’s shape and promoting the intracellular trafficking of key cellular components. Currently, our understanding of these mechanisms remains incomplete. In this study, we identified the non-muscle myosin 10 (MYH10) as an important regulator of adipogenesis and adipocyte function through its interaction with the insulin dependent, Glucose transporter 4 (GLUT4). MYH10 depletion in preadipocytes resulted in impaired adipogenesis, with knockdown cells exhibiting disrupted morphology and reduced molecular adipogenic signals. MYH10 was shown to be in complex with GLUT4 in adipocytes, an interaction regulated by insulin induction. The missing adipogenic capacity of MYH10-KD cells was restored when they uptook GLUT4 vesicles up from neighbor wild-type cells in a co-culture system. Our results provide the first demonstration that MYH10 interacts with GLUT4 in cells and adipose tissue through the insulin pathway. The signaling cascade is regulated by the protein kinase C ζ (PKCζ), which interacts with MYH10 to modify the localization and interaction of both GLUT4 and MYH10 in adipocytes as PKCζ inhibition resulted in reduced GLUT4 and MYH10 translocation and interactions. Overall, our study establishes MYH10 as an essential regulator of GLUT4 translocation, affecting both adipogenesis and adipocyte function, highlighting its importance in future cytoskeleton-based studies in adipocytes.

## Introduction

The process of adipocytes differentiation is termed adipogenesis. It is a well-coordinated process orchestrated by morphological and molecular changes in the cells and their niche. Specific physical, molecular, and chemical pathways promote the commitment of mesenchymal cells to preadipocytes and later to mature adipocytes through a series of sequential inter-dependent events^1,2^. During adipogenesis, the cells undergo substantial morphological changes that are crucial for their commitment to a lineage-specific fate^3,4^. These changes are organized by cytoskeletal components responsible for determining and maintaining cellular shape and function and are a prerequisite for the induction of adipogenic signaling^5–9^. In addition, cytoskeletal proteins are also involved in the terminal differentiation phase and the insulin signaling pathway, where they are needed as a filament-based network for translocation of glucose transporter 4 (GLUT4) and other related processes^10–15^.

GLUT4 is an insulin-dependent glucose transporter expressed mainly in the brain, muscle cells, and adipocytes^16,17^. Reduced GLUT4 translocation and reduced membranal expression, specifically in adipose tissue, is associated with diabetes, insulin resistance, and lowered glucose sensitivity^18,19^. In-vitro, GLUT4 expression is upregulated during adipogenesis, and knockdown of GLUT4 in preadipocytes can interfere with their differentiation^20,21^. GLUT4 is packed in storage vesicles (GLUT4 storage vesicles: GSVs) that are found in the cytoplasm. Upon stimulation, these GSVs are translocated to the cell membrane, where they uptake glucose^16,17^. The translocation process is highly synchronized by a variety of kinases and cytoskeletal proteins that undergo rapid reorganization to promote shuttling of GSVs to the cell membrane ^11,13,22^. Some of the most significant cytoskeletal components associated with GLUT4 translocation are myosins, where different types of myosins interact with both the actin network and the GSVs to facilitate the shuttling process^23–29^.

Non-muscle myosin II (NMII) proteins are vital cytoskeletal components that are ubiquitously expressed in a variety of cell types. The three NMII mammalian paralogs (A-C), encoded by the MYH9, MYH10, and MYH14 genes respectively, differ in their expression profiles, and play both unique and overlapping roles ^30,31^. These proteins interact with actin filaments and are associated with intracellular forces, organelle shuttling, cell adhesion, directional motility, and morphogenesis ^30,32^. The NMII proteins also play a role in the cytoskeletal organization of stress fibers and anchor the cells to the substrate^33,34^. However, despite extensive research addressing the roles of NMII, it remains unclear at the conceptual level how the specific expression profile of MYH10 in individual cells is linked to cell physiology. While a variety of myosin types are implicated in adipocyte metabolism, little is known about the role of MYH10 in adipocytes and adipogenesis. Previous studies have reported that MYH10 is expressed in adipocytes^29,35–38^, and we have previously demonstrated the association of reorganized actin filaments and other candidate proteins, including MYH10, in adipocyte differentiation^6^. However, the mechanism of action and function of MYH10 in adipocytes has never been studied.

Here, we generated a knockdown model of MYH10 in preadipocytes in order to examine its effects on adipogenesis and adipocyte function. The results identify MYH10 as a possible regulator of adipogenesis in 3T3-L1 cells. They also reveal the presence of MYH10 in a complex with GLUT4 that is regulated by insulin. The interaction with GLUT4 was proven crucial for adipogenesis as the lack of adipogenic capacity of MYH10 knockdown cells was restored by transportation of GLUT4 from neighboring cells. Moreover, our results indicate that the MYH10:GLUT4 complex is regulated by PKCζ, a protein kinase C associated with insulin signaling and cytoskeleton reorganization^39,40^. This is the first study to show the importance of MYH10 in adipogenesis and adipocyte function and can serve as the foundation for future MYH10 based studies in adipocytes.

## Methods and Materials

### Animals

Epididymal visceral adipose tissues were taken from C57bl/6J mice and used as fresh and frozen tissues for further procedures. The mice were kept in a conventional facility with 12 h light/dark cycles and were fed with standard chow and provided water ad libitum. Animal care and experiments were in accordance with the guidelines of the IACUC Approval (01-21-044).

### Cell lines

Mouse embryonic 3T3-L1 preadipocytes (American Type Culture Collection) were cultured and differentiated as was previously described^41^. For insulin induction, differentiated 3T3-L1 cells were incubated in a Dulbecco’s modified Eagle’s medium (DMEM) without glucose for one hour (Biological Industries, Israel), and then replaced with a GM with and without 5µg/ml insulin for 5-30 minutes. To inhibit PKCζ activity, a myristoylated pseudo-substrate inhibitor for PKCζ (Santa Cruz, SC -397537) was added at 50µM to the starvation and induction phases of insulin induction.

### Lentivirus production and transduction

Lentiviruses were produced as was previously described^42^. The pLenti-myc-GLUT4-mCherry expressing lentivirus plasmid was a gift from Weiping Han (Addgene plasmid # 64049; http://n2t.net/addgene:64049; RRID: Addgene_64049)^43^. The MYH10 pCMV-GFPH2B lentiviral plasmid (Clone ID; TRCN0000110555) was a gift from Chen Luxenburg, Tel Aviv University, Tel Aviv, Israel.^44^ The lentivirus particles were produced by the transfection of cultured HEK293FT cells (Invitrogen; R70007) with the lentivirus expression plasmid and helper plasmids [pLP1, pLP2, and VSV-g (Invitrogen)]. The supernatant was collected after one and 2-days post-transfection. Cultured 3T3-L1 preadipocytes were infected control (Scr; H2B-GFP) or gene-specific lentiviruses with Polybrene (Sigma-Aldrich) at a final concentration of 100µg/ml.

### Co-culturing

For the GLUT4^+^: MYH10-KD 3T3-L1 co-culture, GLUT4^+^ cells were seeded first, and after one day, the MYH10-KD cells were added to all wells. The culture was then differentiated as described. The PKH26:MYH10-KD co-cultures 3T3-L1 cells were labeled with 0.5λ PKH26 Fluorescent Cell Linker Kit (Sigma-Aldrich); the GFP cells were added after one day.

### Immunofluorescence staining

Immunofluorescence staining was performed as described in Mor-Yosef Moldovan et al^6^. Shortly, the cells were fixed with a 4% paraformaldehyde solution, permeabilized with 0.5% Triton in 1% TBST, and then blocked with a blocking solution (1% TBST containing 1–2% normal goat serum and 1% BSA). Next, the cells were incubated overnight with primary MYH10 (Santa Cruz; SC-376942) and GLUT4 (Santa Cruz; SC-53566) antibodies, washed, and incubated with secondary antibodies, Cy3-anti-mouse (115-165-003; Jackson ImmunoResearch Laboratories), Alexa Fluor 555 anti-Mouse IgG1 (Invitrogen; A-21127), and Alexa Fluor 488 anti-Mouse IgG2b (Invitrogen; A-21141) for one additional hour. F-actin filaments were stained with fluorescein isothiocyanate labeled phalloidin (P5282; Phalloidin-FITC; Sigma-Aldrich). The stained coverslips were mounted on slides with Fluoroshield™ mounting medium containing 4’, 6-diamidino-2-phenylindole (DAPI). Images were acquired by a confocal microscope (Leica SP8; Leica, Wetzlar, Germany) and a fluorescence microscope (Nikon, Eclipse Ci).

### Whole-mount staining

Adipose tissue whole-mount staining was performed as previously described^45^. Briefly, isolated murine epididymal adipose tissues were fixated in 1% paraformaldehyde in 24-wells plates. The tissues were then washed and blocked with a blocking buffer (PBS-0.3T with 5% normal goat serum). Blocked tissues were incubated overnight with primary MYH10 (Santa Cruz; SC-376942) and GLUT4 (Santa Cruz; SC-53566) antibodies. Next, the tissues were incubated with secondary antibodies, Alexa Fluor 555 anti-Mouse IgG1 (Invitrogen; A-21127), and Alexa Fluor 488 anti-Mouse IgG2b (Invitrogen; A-21141) and washed again before adding the Fluoroshield™ mounting medium DAPI. Images were acquired by a confocal microscope (Leica SP8; Leica, Wetzlar, Germany).

### Image processing and analysis tools

ImageJ was used to analyze and process the immunofluorescence pictures. ***Cytoskeleton quantification***: MYH10 filaments distribution analysis was done as previously described^6^. Shortly, the images were analyzed with the FIJI ImageJ software (NIH, Bethesda, MD) using two plugins. OrientationJ^46^ was used to quantify the coherency of the cells, and the Ridge detection plugin^47,48^ was used to calculate the length and number of junctions of each cell. ***Membranal/cytoplasmatic ratio quantification***: Membranal/cytoplasmatic and cortical/cytoplasmatic ratios were calculated as previously described^49^. The mean membranal and cytoplasmatic intensities were extracted by using the method. The mean membranal intensity was divided by the mean membranal intensity to extract the ratio of cells.

### Live microscopy

All live imaging was performed using EVOS FL Auto 2 microscope (Invitrogen). ***Adhesion assay*:** Suspended 3T3-L1 cells were seeded in six wells and were immediately transferred to the EVOS microscope. Phase-contrast images were taken at 3, 15- and 33 minutes post-seeding. Fiji ImageJ’s Trainable Weka segmentation plugin^50^ was used to separate the cells and background in each image. Every cell in each field was marked, and its area was calculated using ImageJ. ***Migration assay*:** The migration of cells was assessed by manual track of single cells using time-lapse images. The cells were observed for three hours, and images were taken at ten minutes intervals. Trajectories of the migration paths were calculated using the manual cell tracker plugin in ImageJ. Each cell nucleus in the image sequence was manually marked for each frame. Then the created trajectories were used to generate the vectors and calculate the motility data. Accumulative distance and average speed were calculated for each trajectory. ***Wound healing assay:*** Cultured confluent (90%) monolayer of 3T3-L1 preadipocytes were scratched with a tip. The cells were then observed under the EVOS microscope and measured at 0, 4, 8, and 12 hours and at the time of closure. The relative closure gap was calculated as the ratio of the current gap relative to the gap at the starting point. The experiment was repeated three times for each group.

### Adipogenesis assays

Images of cultured 3T3-L1 cells throughout differentiation were taken in order to follow the cell’s growth, morphology and LDs accumulation. ***Level of adipogenesis:*** The Level of adipogenesis was calculated as previously described^51^. Shortly, Stitched phase-contrast x40 images of differentiated cultures were taken after 21 days post-adipogenesis induction. Based on the major visual difference between fibroblasts and adipocytes a visual difference mapping (VDM) was obtained. The map was used to calculate the level of adipogenesis (LOA) in each culture. ***Lipid droplet quantification and morphological analysis*:** Phase contrast Images at a magnification of x400 of differentiating cultures were taken 21 and 28 days post-induction. The LDs radius and cell projected area were analyzed using an image-processing-based method developed by Lustig et al^51^.

### Immunoblotting

The procedures and analyses were performed according to the standard protocols (www.protocol-online.net). Cells were harvested from cultures, washed with ice-cold PBS, and lysed in 50 mM Tris pH 7.5, 150 mM NaCl buffer containing 1 mM EDTA, 1% NP-40 and protease inhibitors: [phenylmethylsulfonyl fluoride (PMSF), 1 mM; 1-chloro-3-tosylamido-4-phenyl-2-butanone, TPCK, 10 µg/ml; aprotinin, 10 µg/ml (Sigma-Aldrich)]. Protein concentration was determined with BCA Protein Assay Kit (Pierce 23225). Samples were re-suspended in Laemmli buffer, separated on 7.5% SDS– PAGE gel, and transferred to nitrocellulose. After blocking the membranes were incubated overnight with a primary antibody, anti–MYH10 (Santa Cruz; SC-376942). Primary antibody was washed 4 times for 5 min with TBST and followed by incubation with Peroxidase Anti-Mouse IgG (Jackson Immuno Research) in blocking solution. Peroxidase signal was detected with chemiluminescent substrate (Pierce, Rockford, IL) using Fusion FX7 (Vilber Lourmat). Coimmunoprecipitation Whole-cell lysates (200µg) were incubated with Protein A/G Plus Agarose (Santa Cruz, SC-2003) and an anti-GLUT4 (SC-53566) antibody overnight. The samples were washed and dissolved in Laemmli buffer.

### Mass spectrometry

Mass spectrometry data was analyzed based on our data base that was previously described^6^

### RNA isolation and qPCR

Total RNA was extracted from 3T3-L1 cells (EZ RNA kit, Biological Industries, Beit Haemek, Israel) and reverse transcribed to cDNA using High-Capacity cDNA Reverse Transcription Kit (Applied biosystems). Transcripts levels were measured with SYBR green (Applied biosystems) using STEPONE plus system (Life Technologies). All data was normalized to Actin by the delta delta CT method^52^.

The sequences of the primers are listed below:

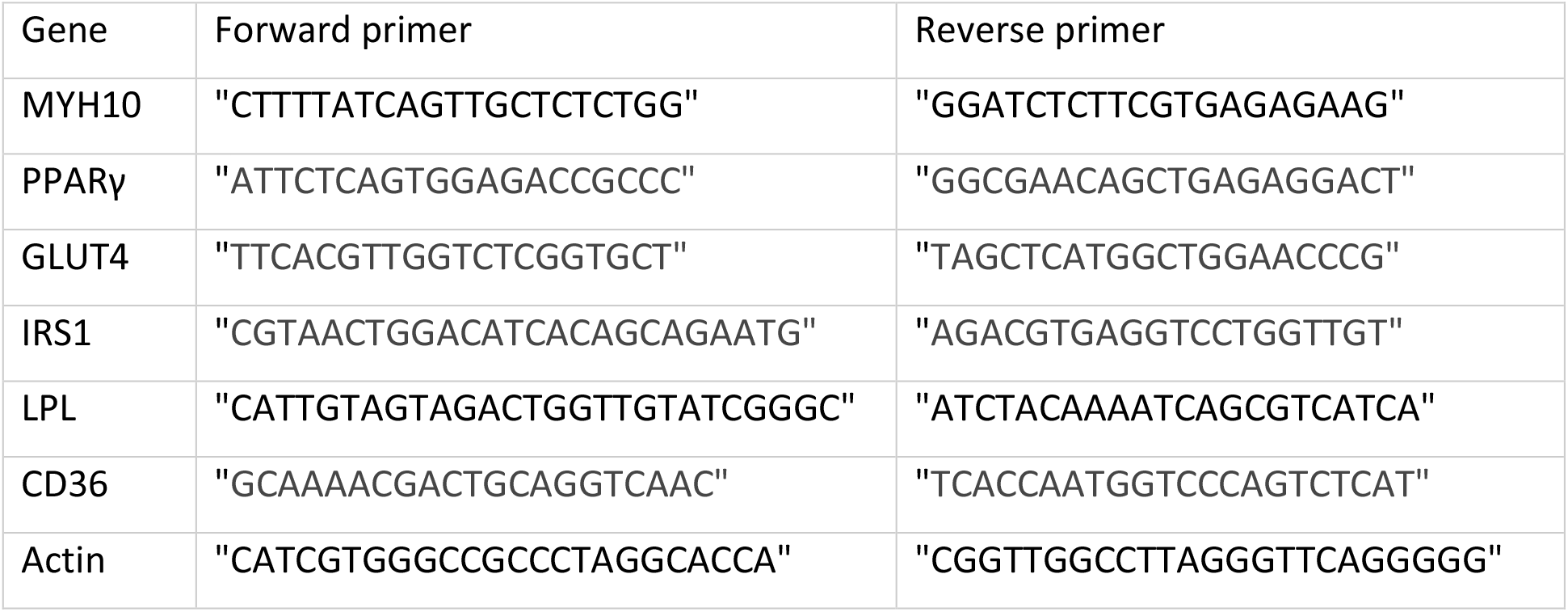

### Bioinformatics pathway analysis

Bioinformatics analysis for the mass spectrometry data was performed using Perseus software as described in Mor-Yossef Moldovan et al^6^. Phosphorylation partners analysis was performed using the PathwatNet website^53^. The top fifty candidate genes were extracted and compared with Insulin pathway signaling pathway proteins (GO:0008286) that were extracted from the gene ontology database^54–56^.

### Statistical analysis

Statistical analyses were analyzed by GraphPad Prism v.8.1.1. Results are presented as means ± SEM. All results were tested for normal distribution by Kolmogorov-Smirnov test, and outliers were identified using the ROUT method. Statistical differences comparing the mean values were tested using two-tailed, unpaired t-tests or one way-ANOVA where appropriate. Values that were not normally distributed were tested using Mann-Whitney or Kruskal-Wallis (for three or more groups), followed by Dunn’s post-test for multiple comparisons. A value of p < 0.05 was considered statistically significant.

<u>Schematic illustrations:</u> were created by Bio Render software https://biorender.com

## Results

### MYH10 distribution is altered during adipogenesis in 3T3-L1 cells

A previous study from our lab examined the association between changes in cell morphology and the reorganization of the actin cytoskeleton^6^. As part of the analysis, we examined the distribution of actin fibers during adipogenesis and identified the non-muscle myosin isoform (MYH10) as a potential cytoskeletal protein that may impact the function of adipocytes which led us to examine the role and function of MYH10 in adipocytes. Immunostaining of MYH10 (red) and F-actin (green) in undifferentiated 3T3-L1 cells showed a colocalization pattern of the filaments as part of the actomyosin cytoskeleton, indicating a mutual role in undifferentiated (UD-F) cells (Fig. 1A). In order to examine whether MYH10 filaments undergo reorganization during adipogenesis in UD-F cells, we compared the distribution of MYH10 in UD-F and differentiated cells (Diff). The results indicated a substantial difference in the binary images of MYH10 during differentiation, where the organized filamentous-like form distributed evenly throughout the cell in undifferentiated cells was disrupted and predominantly located in a cortical ring around the cell membrane once the cells differentiated (Fig. 1B). This prompted us to examine the dynamic nature of MYH10 reorganization during the differentiation of adipocytes. Comparison of MYH10 in cells at different stages in the same culture revealed that the coherency in UD-F was significantly higher than in Diff (0.14 and 0.016, respectively), with more junctions than in Diff cells. Moreover, the MYH10 filaments in UD-F were 1.7 times longer in the undifferentiated cells (Fig.1C). These observations suggest that, like actin, MYH10 undergoes significant reorganizations during adipogenesis.

**Fig.1.**
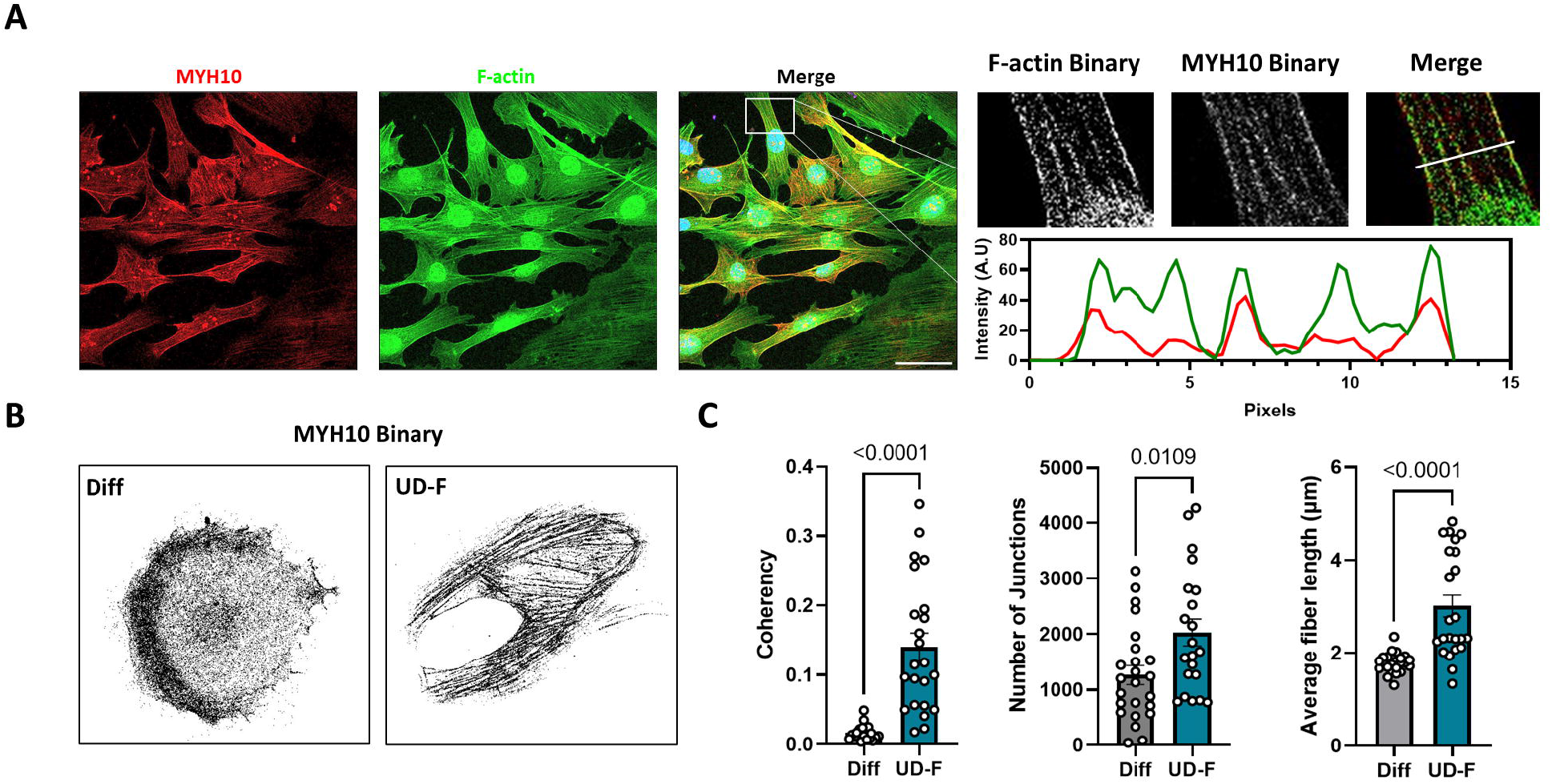
MYH10 distribution is altered during adipogenesis in 3T3-L1 cells. (A) Immunostaining of preadipocytes (magnification of X630, scale bar=50µm) stained for MYH10 (red), Phalloidin (green) and DAPI (blue) and an enlargement of MYH10 and actin filament colocalization with a line scan of corresponding fluorescence intensities of MYH10 (red) and actin (green) (B) A binary illustration of MYH10 staining in an adipocyte (Diff) and a preadipocyte (UF-D) (C) Quantification of the coherency, number of junctions and length of MYH10 in adipocytes (Diff, n=24) and preadipocytes (UF-D, n=24), Significance was calculated using unpaired nonparametric Mann-Whitney test. Error bars represent means ± SEM.

### MYH10 knockdown model

As the next stage in exploring the effect of MYH10 on adipocyte function, we generated an shRNA-MYH10 knockdown system in 3T3-L1 cells infected with an shRNA-MYH10 lentivirus containing a histone 2B green fluorescent protein (GFP) marker (MYH10-KD). The knockdown efficacy was tested by qPCR, which confirmed a substantial decrease in MYH10 levels (Fig.2A). Knockdown efficacy was also examined by analyzing the percentage of transfected cells (GFP^+^) and examining the intensity of cytoplasmatic MYH10 in the GFP^+^ cells. The results indicated that 80% of cells in the culture were GFP positive, and that these cells exhibited lower levels of cytoplasmatic MYH10 (Fig. 2B).

**Fig. 2.**
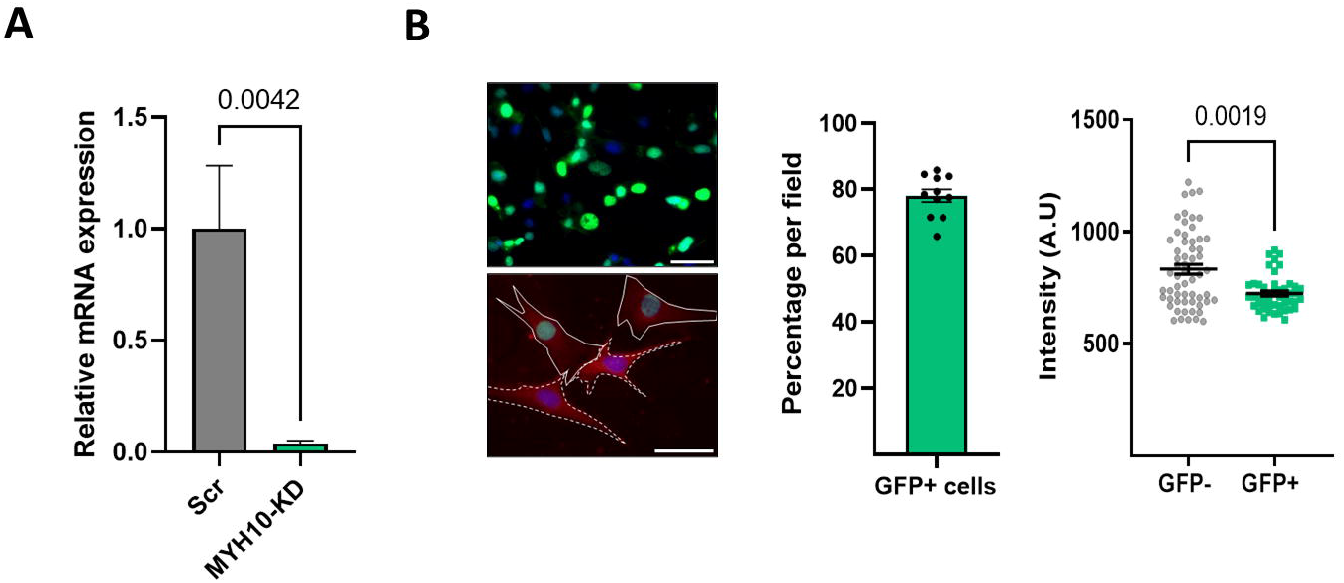
MYH10 KD model. (A) mRNA expression of MYH10 in Scramble and MYH10-KD cells, measured in preadipocytes. (B) Immunostaining of DAPI (blue) and MYH10-H2B-GFP (green nuclei) in MYH10-KD cultures (magnification of X400, scale bar=50 µm), Immunostaining for MYH10 (red), DAPI (blue), and MYH10-H2B-GFP (green nuclei) in MYH10-KD preadipocyte cultures (scale bar equal 50 µm), morphometric analysis of the average GFP positive cells in MYH10-KD cultures and a single-cell analysis of MYH10 fluorescent intensity in the cytoplasm, significance was calculated using an unpaired nonparametric Mann-Whitney test, error bars represent means ± SEM.

### MYH10 KD affects cell motility and migration

Since MYH10 is known to play roles in cell migration and adhesion, we assessed the effect of its depletion on cell adhesion and motility in knockdown (KD) 3T3-L1 preadipocytes and cells infected with a scrambled construct (scrambled). Figure 3A presents the differences in adhesiveness between the cultures. After 33 minutes of adhesion, the cell projected area of control scrambled cells was 2.5-fold higher than for KD cells, suggesting a possible effect of MYH10 on cell spreading and adhesion in preadipocytes (Fig. 3B). Next, we examined the impact of MYH10 KD on cell migration. The results of a wound-healing assay (illustrated in fig. 3D), with the relative scratch gap measured after 4, 8, and 12 hours, revealed a significantly larger gap at the endpoint in MYH10 KD cells than scrambled cells (40% vs. 8% respectively, Fig. 3C, D). With respect to migration, the average speed and accumulated distance of individual KD cells were reduced by more than 20%, demonstrating the effect of MYH10 on the motility of the cells (Fig. 3E,F). Figure 3G presents representative images of the trajectories of cells after 1 and 3 hours. These data support the suggestion that MYH10 is a key factor in adhesion and migration.

**Fig. 3.**
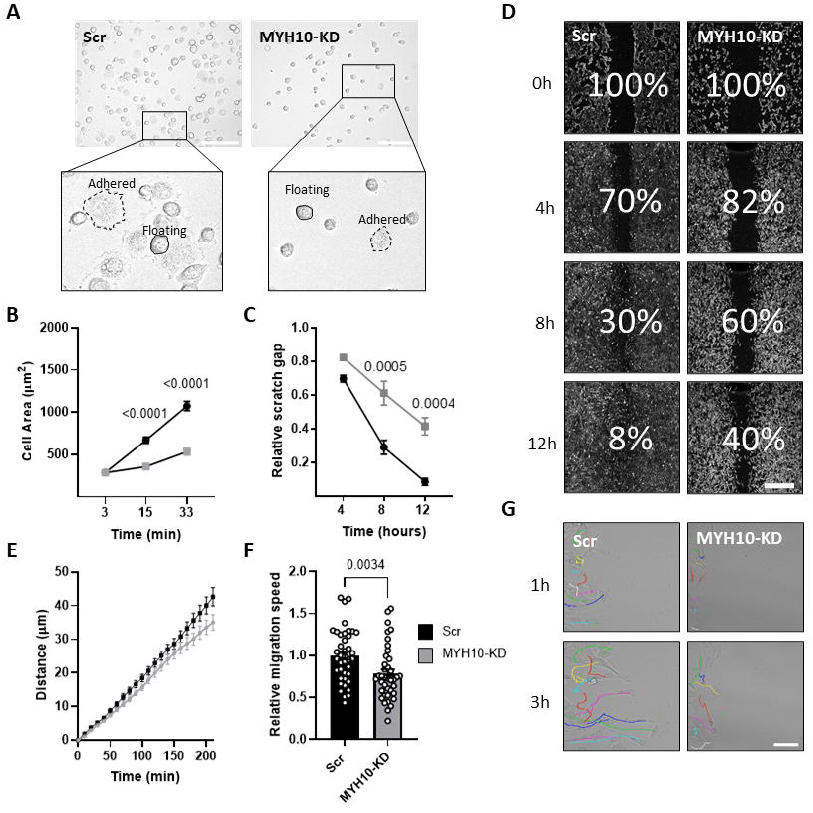
MYH10 KD affects cell motility and migration. (A) Representative phase-contrast images from an adhesion assay of Scramble (Scr) and MYH10-KD cells 21 minutes after they were seeded, demonstrating the differences between Scr and MYH10-KD cells’ adhesion rates. (Magnification X200, scale bar, 125µm). (B) Single-cell analysis of the mean cell area at the indicated time points (n>90). Significance was calculated using Two-way ANOVA with Sidak’s post-test. (C) Wound healing assay, the ratio of the remaining gap at a given time to the gap at the starting point, Scramble (black) and KD cells (grey) (n=3), Significance was calculated using Two-way ANOVA with Sidak’s post-test. (D) Representative pictures of a wound-healing assay of the Scr and MYH10-KD cells at the indicated time points (magnification of X40, scale bar equals 650μm) (E) Illustrative Images of the migration trails of Scr and MYH10-KD cells at the indicated time points. Each colored path illustrates the migration of an individual cell over time (magnification X200, scale bar, 125 µm). (E) Accumulative distance and (F) mean speed of Scramble and MYH10-KD cells (n=40), Significance was calculated using unpaired nonparametric Mann-Whitney test. (G) Illustrative Images of the migration trails of Scr and MYH10-KD cells at the indicated time points. Each colored path illustrates the migration of an individual cell over time (magnification X200, scale bar, 125 µm). Error bars represent means ± SEM.

### MYH10 knockdown affects adipogenesis

In order to assess the effect of MYH10 on adipogenesis, scrambled and KD cultures were differentiated to adipocytes and the level of adipogenesis was evaluated after 21 and 28 days. As can be seen in Figure 4A, C, knockdown cultures displayed a lower LOA than the scrambled cultures; KD cells exhibited little to no adipogenesis, and the fraction of differentiated cells were primarily due to the WT (Wild type; GFP^-^) cell population in the culture. Quantification of the LOA (the percentage of adipocytes in the culture relative to the scrambled cells) confirmed the reduction in adipogenesis in the knockdown cells (Fig. 4B). With respect to adipogenesis-related morphologic parameters, scrambled cells had a significantly greater cell area than the MYH10 KD cells (1800 μm^2^ versus 520 µm^2^, respectively). They also had larger lipid droplets (3.3 μm in the scrambled cells compared to 1.5 μm, respectively (Fig. 4E)). Both the cell area and size of lipid droplets rose significantly between day 21 and day 28 of differentiation, but only in the scrambled cells, suggesting that adipogenesis ceased mid-differentiation in the KD cultures. Figure 4F presents the reduction in the expression of principal adipogenic markers in the knockdown cultures, with significant downregulation of all measured markers (PPARγ, GLUT4, IRS1, LPL, and CD36). This observation supports the results of a lack of adipogenesis due to the knockdown of MYH10, and together, these data strongly suggest a major role for MYH10 in adipocyte differentiation.

**Fig. 4.**
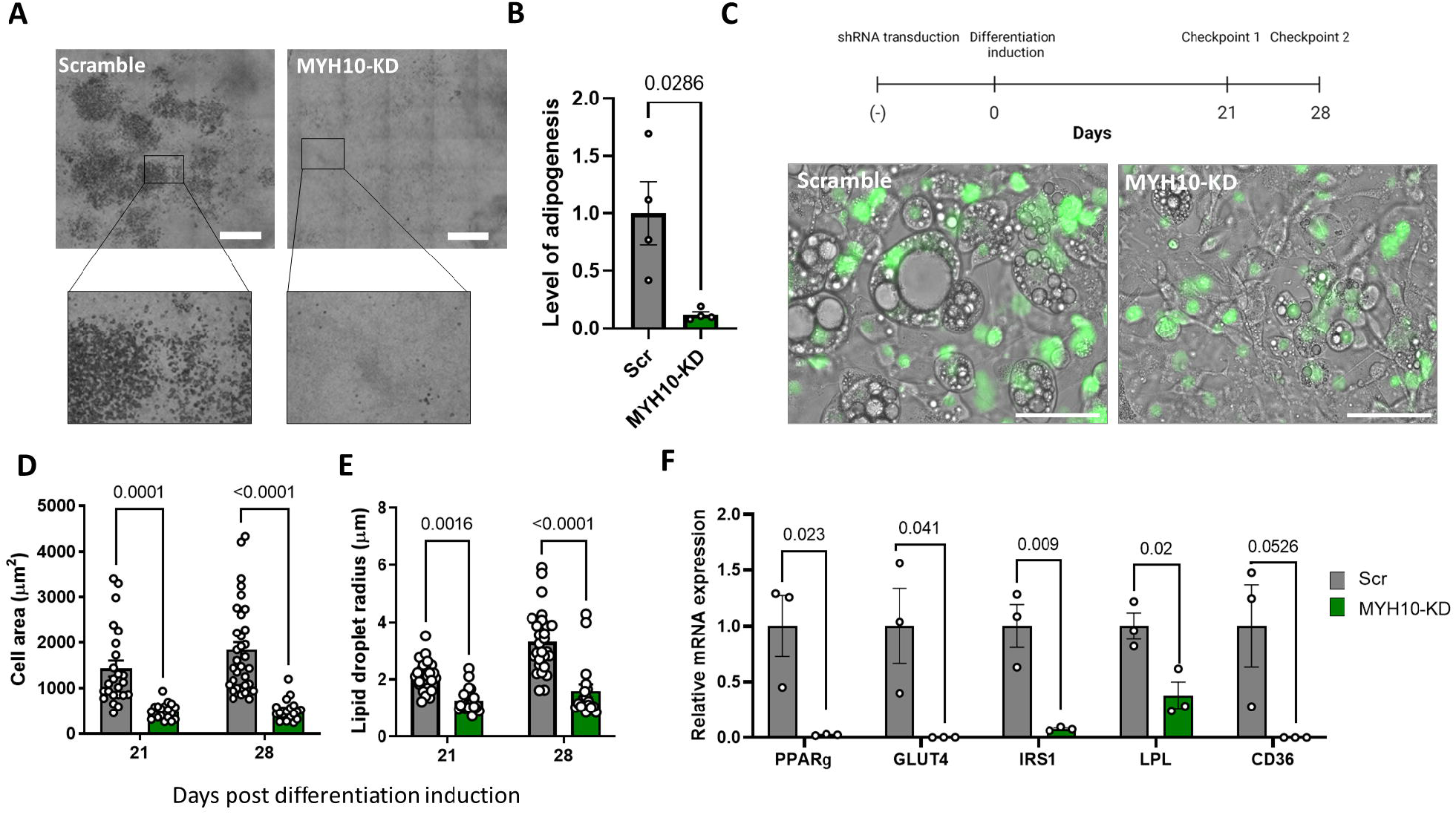
MYH10 knockdown affects adipogenesis. (A) Images of scrambled (Scr), and MYH10-KD cultures, at 28 days after initiation of adipogenesis (scale bar=650 µm) (B) level of adipogenesis (percentage of adipocytes in the culture) of Scr (grey, n=4) and MYH10-KD (green, n=4) cultures, day 28 post differentiation, significance was calculated using unpaired nonparametric Mann-Whitney test. (C) The experimental model and phase contrast overlay with florescent GFP (H2B-GFP) at 28 days after initiation of adipogenesis in Scr and MYH10-KD cells (scale bar equals 75 µm, magnification of X400). Morphology measurements for (E) cell-projected area and (E) lipid droplets radius for Scr, and MYH10-KD cells, measured 21 and 28 days after adipogenic induction, significance was calculated using one-way ANOVA with Tukey’s post-test (p<0.0001) (F) quantitative PCR of Scr (grey) and MYH10-KD (green) cells (n=3 per group) for PPARγ, GLUT4, IRS1, LPL and CD36, significance was calculated by an unpaired student t-test profile. Error bars represent means ± SEM.

### Co-expression of glucose transporter 4 and MYH10

While MYH9, an MYH10 paralog, is known to interact with GLUT4 and was shown to affect its intracellular translocation, this aspect of MYH10 has never been extensively studied. Immunostaining of MYH10 and GLUT4 in murine visceral adipose tissue revealed a similar cortical/membranal expression pattern in mature adipocytes (Fig. 5A). Co-immunoprecipitation of adipose tissue preparations with GLUT4 followed by a western blot with MYH10 demonstrated an association between the two proteins and suggested the presence of a GLUT4:MYH10 complex in the tissue (Fig. 5B). As for adipose tissue, immunofluorescence staining in 3T3-L1 cells demonstrated colocalization of MYH10 and GLUT4, particularly in the cell membrane and the cortical area of the cells (Fig. 5C). Similarly, GLUT4 and MYH10 also co-immunoprecipitated from differentiated 3T3-L1 cells, indicating that the proteins exist together in a complex both in-vivo and in-vitro (Fig. 5D). Mass spectrometry of undifferentiated and differentiated adipocytes revealed the expected differential expression of glucose transporters. GLUT1 is expressed in fibroblasts, and the level increases in adipocytes, while GLUT4 is expressed only by differentiated adipocytes, highlighting the importance of GLUT4 in adipogenesis (Fig. 5E). These data suggest that the effect of MYH10 on adipogenesis may be related to the interaction with GLUT4.

**Fig. 5.**
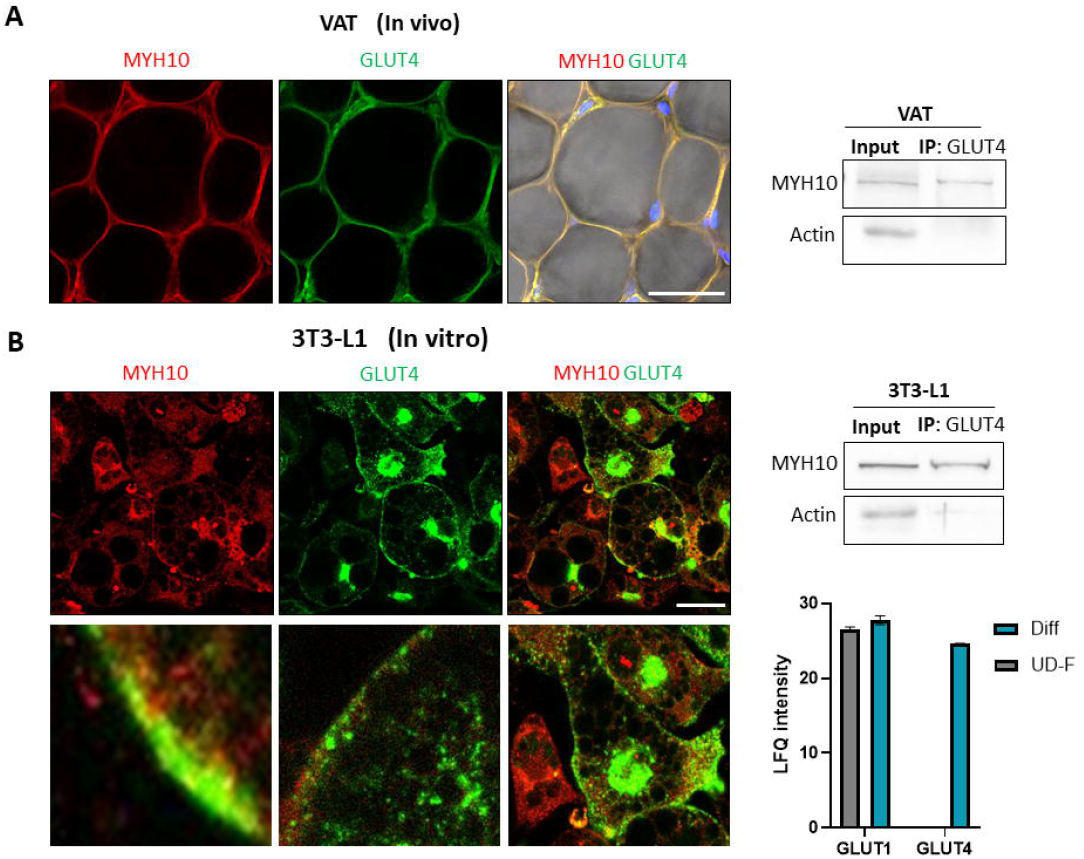
Co-expression of Glucose transporters 4 and MYH10. (A) Whole mount staining of MYH10 (red), GLUT4 (green) and DAPI (blue) in murine visceral adipose tissue (Magnification of X200, scale bar=50μm) and Co-Immunoprecipitation of GLUT4 and WB for MYH10 and Actin in murine visceral adipose tissue (B) Immunofluorescence staining of MYH10 (red), GLUT4 (green) and DAPI (blue) in adipocytes. (Magnification of X630, scale bar=50μm), Co-Immunoprecipitation of GLUT4 and WB for MYH10 and Actin in differentiated 3T3-L1 cells and a LFQ intensity histogram from a mass spectrometry analysis for GLUT4 and GLUT1 in undifferentiated (Diff) and differentiated 3T3-L1 cells.

### MYH10 colocalization and interaction with GLUT4 is induced by insulin

Since GLUT4 translocation is insulin-dependent, we examined the changes in the expression and localization of MYH10 in response to insulin. After differentiation, 3T3-L1 cells were starved for one hour and then induced with insulin for a further 30 minutes. Immunostaining of MYH10 and GLUT4 in induced and uninduced cells revealed that insulin-induction strongly increases the expression of GLUT4 in the membranal compartment. MYH10 exhibited a similar pattern, with prominent expression in the cortical region of stimulated cells (Fig. 6A). Intensity plots also revealed colocalization of MYH10 and GLUT4 after induction (Fig. 6B). The membranal to cytoplasmatic ratio (MCR) of GLUT4 was 1.5-fold higher after insulin stimuli, meaning that more GLUT4 was translocated to the membrane post-induction (Fig. 6C). Quantification of MYH10 MCR also demonstrated a similar increase post stimulus (1.48-fold, Fig. 6C). We then examined the effect of insulin induction on total MYH10 expression; a western blot of MYH10 after 15 and 30 minutes showed no difference in MYH10 levels indicating that insulin primarily affects the localization and function but not the expression levels of MYH10 in adipocytes (Fig 6D). Co-immunoprecipitation with GLUT4 followed by a western blot with MYH10 revealed an increase in the MYH10:GLUT4 complex in cells after insulin induction, implying a functional role for MYH10 in GLUT4 translocation in induced adipocytes (Fig. 6E). Overall, these results indicate that MYH10 is connected to the insulin pathway with a possible effect on GLUT4 translocation from the cytoplasm to the cell’s membrane.

**Fig. 6.**
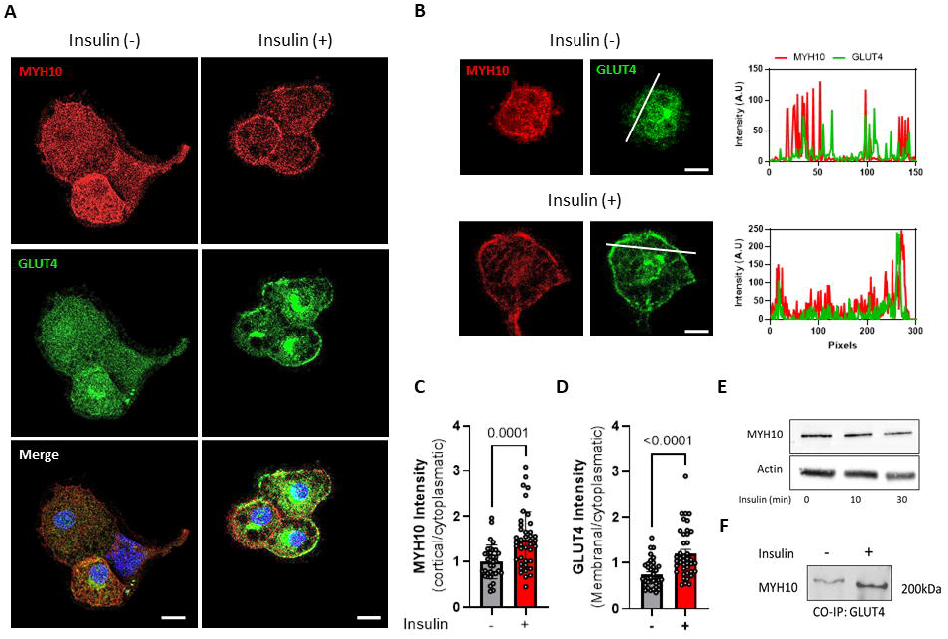
MYH10 localization and interaction with GLUT4 is induced by insulin. (A) Immunofluorescence staining of MYH10 (red), GLUT4 (green) and DAPI (blue) in differentiated 3T3-L1 cells +/- 30 min of insulin induction. (Magnification of X630, scale bar=50μm). (B) Enlargements of differentiated 3T3-L1 cells (+/- insulin) and Intensity line profiles of MYH10 (red) and GLUT4 (green) of the membranal and cytoplasmatic profile. (Magnification of X630, scale bar =50 μm). (C) Cortical to cytoplasmatic intensity ratio quantification of MYH10 in differentiated insulin induced (n=41) and non-induced (n=37) 3T3-L1 cells, significance was calculated using unpaired nonparametric Mann-Whitney test. (D) Membranal to cytoplasmatic intensity ratio quantification of GLUT4 in differentiated insulin induced (n=41) and non-induced (n=37) 3T3-L1 cells, significance was calculated using unpaired nonparametric Mann-Whitney test. (E) Western blot analysis of MYH10 in differentiated 3T3-L1 cells 0,15- and 30-minutes post insulin induction (F) Co-Immunoprecipitation of GLUT4 and WB for MYH10 in differentiated 3T3-L1 cells +/- 30 min of insulin induction. Error bars represent means ± SEM.

### GLUT4^+^ shuttling can restore the adipogenic capacity of MYH10 KD cells

In order to determine whether the adipogenic capacity of MYH10 cells can be restored by the transportation of GLUT4 from neighboring cells, we established a co-culture model of 3T3-L1 cells expressing a GLUT4-mCherry (GLUT4^+^) with MYH10-KD cells expressing nuclear GFP. The labeled GLUT4 (red) in GLUT4^+^ enables us to track the movement of GLUT4 between cells and analyze cell-to-cell interactions. Co-cultures of GFP Scramble cells with the preadipocytes stained with PKH26 red fluorescent dye or marked GLUT4^+^ cells were used to examine the basic cell-to-cell interaction model. The results presented in Fig. 7A clearly show the uptake of red-labeled particles by the GFP^+^ cells in both cases. Phase-contrast images of differentiated GLUT4^+^ and MYH10-KD monocultures demonstrated their disparate adipogenic capacity (Fig. 7B). Images of the differentiated co-cultures revealed numerous GFP^+^ adipocytes filled with GLUT4-mCherry staining (Fig. 7C). Unsurprisingly, the LOA of co-cultures of MYH10-KD and GLUT4^+^ was higher than GFP^+^ cells alone (Fig. 7D). Strikingly, the number of differentiated GFP^+^ cells per field was significantly higher in the co-culture (average of 4 compared to < 1 in the GFP culture, Fig. 7E). These results may suggest that labeled GLUT4 is taken up by GFP^+^ cells to compensate for the lack of GLUT4 transferred from within the cell to its outer membrane and indicate the presence of cellular communication that can transfer GLUT4 from neighboring cells to support and induce the differentiation of MYH10-KD cells.

**Fig. 7.**
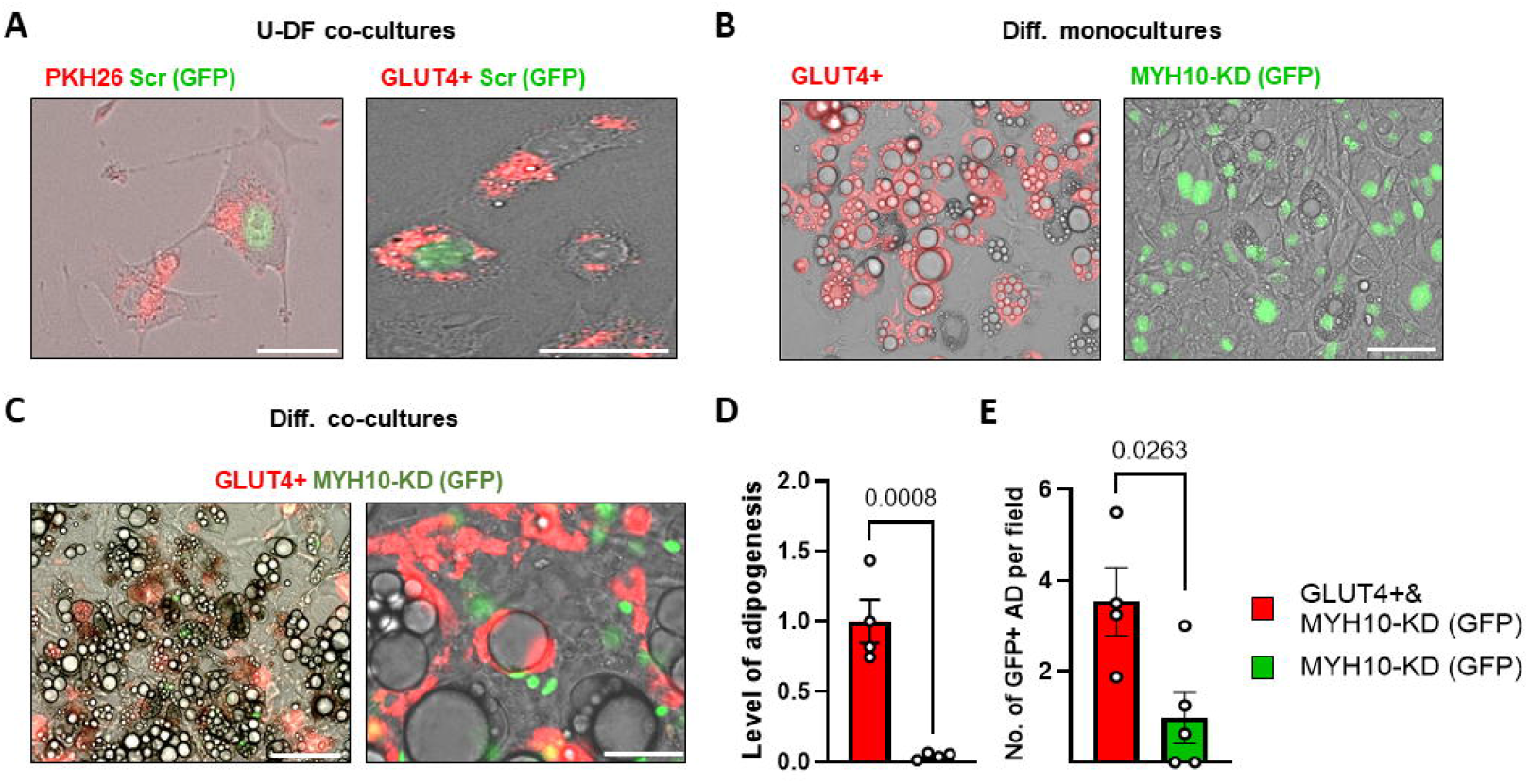
GLUT4^+^ cells can restore the adipogenic capacity of MYH10 KD cells. (A) Image of vesicles stained with PKH26 (right) that were internalized by GFP^+^ cells (green nucleus) and image of differentiating MYH10-KD (green nucleus) cells with internalized GLUT4-mcherry (red) (scale bar 62.5μm) (down). (B) Phase contrast overlay with florescent GLUT4^+^ (left) and MYH10-KD (right) differentiated cultures (scale bar=125µm, magnification of X400). (C) Image of MYH10-KD (green nuclei) in a co-culture with GLUT4^+^ Cells 28 days post differentiation. (Scale bar=125 (right) and 75µm (left), magnification of X400). (D) level of adipogenesis (percentage of adipocytes in the culture) of co-cultures of GLUT4 cells with MYH10 KD cells (light grey) and MYH10-KD cells (dark grey), significance was calculated using a two-tailed unpaired student’s t-test. (E) Number of differentiated GFP^+^ cells in a culture in co-cultures of GLUT4^+^ cells with MYH10-KD cells (light grey) and MYH10-KD cells (dark grey), significance was calculated using a two-tailed unpaired student’s t-test, error bars represent means ± SEM.

### MYH10 interaction with GLUT4 post insulin induction is mediated by PKCζ

A number of pathways and effectors are known to regulate the function of MYH10 through phosphorylation. We used PathwayNet analysis, a web-based tool for diverse protein-to-protein interactions to identify potential phosphorylation partners of MYH10 in order to better understand its involvement in insulin-induced GLUT4 translocation (Fig. 8A). After retrieving the top PathwayNet phosphorylation candidates, we crossmatched them to the gene ontology list of insulin signaling pathway-related proteins. PKCζ, a protein kinase C protein, was the lead candidate, since it is a known component of the insulin signaling pathway and a potential MYH10 phosphorylation partner. In order to further explore the hypothesis that PKCζ affects MYH10 function, we used a myristoylated pseudo-substrate inhibitor for PKCζ to examine the effect on MYH10 activity. Insulin-induced differentiated 3T3-L1 cells were incubated with and without the inhibitor to examine its effect on MYH10 and GLUT4. As shown in Fig. 8B-C, insulin induction triggered a ring-like membranal expression of both MYH10 and GLUT4. In contrast, the inhibitor dramatically disturbed the translocation and localization of MYH10 and GLUT4, resulting in chaotic and disruptive expression of both proteins.

**Fig. 8.**
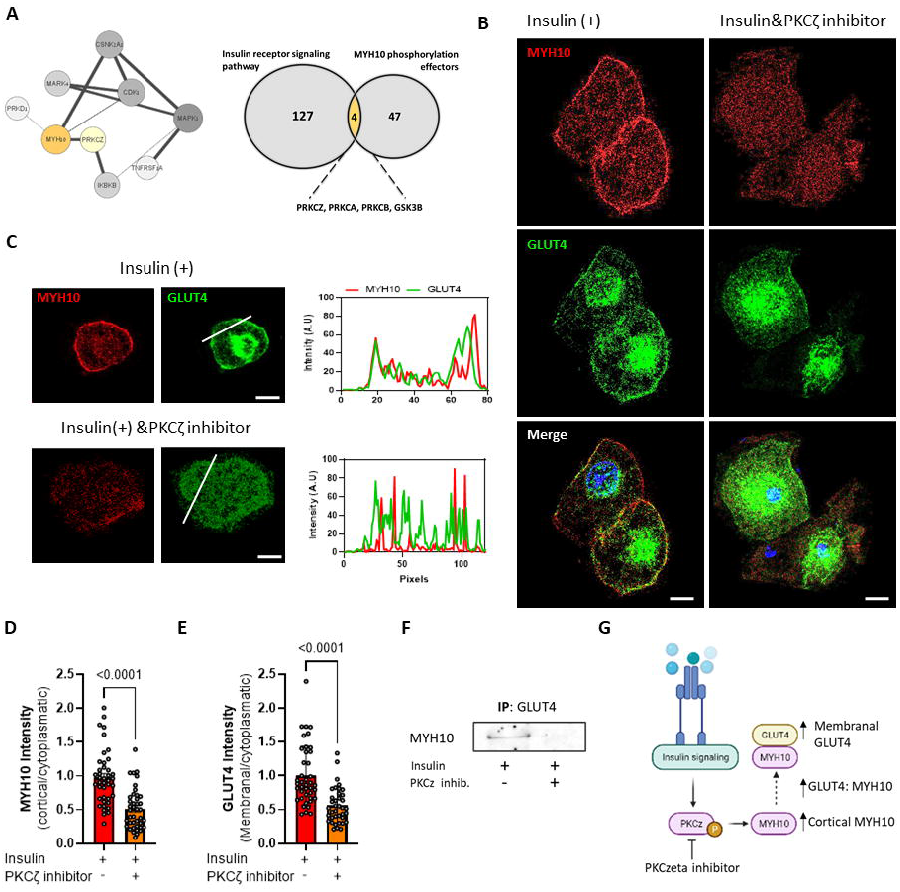
MYH10 interaction with GLUT4 post insulin induction is mediated by PKCζ. (A) A network analysis of MYH10 potential upstream phosphorylation partners. Generated by PathwayNet and a Venn diagram of GO:0008286, insulin receptor signaling pathway proteins and MYH10 potential upstream phosphorylation partners intersection. (B) Immunofluorescence staining of MYH10 (red), GLUT4 (green) and DAPI (blue) in differentiated 3T3-L1 cells after 30 min of insulin induction w/o PKCζ pseudosubstrate inhibitor. (Magnification of X630, scale bar=50μm). (C) Enlargements of differentiated 3T3-L1 cells +/- the PKCζ pseudosubstrate inhibitor and Intensity line profiles of MYH10 (red) and GLUT4 (green) of the membranal and cytoplasmatic profile. (Magnification of X630, scale bar=50μm). (D) Quantification of the ratio between the cortical and cytoplasmatic intensity of MYH10 in differentiated insulin induced 3T3-L1 cells with PKCζ inhibitor (n=41, orange) and without PKCζ inhibitor (n=41, red), significance was calculated using unpaired nonparametric Mann-Whitney test. (E) Quantification of the ratio between the Membranal and cytoplasmatic intensity of GLUT4 in differentiated insulin induced 3T3-L1 cells with PKCζ inhibitor (n=41, orange) and without PKCζ inhibitor (n=41, red), significance was calculated using unpaired nonparametric Mann-Whitney test (F) Co-Immunoprecipitation of GLUT4 and WB for MYH10 in differentiated 3T3-L1 cells after 30 min of insulin induction w/o PKCζ inhibitor. (G) Schematic illustration of the purposed role of PKCζ and MYH10 in insulin signaling. Error bars represent means ± SEM.

Furthermore, MCR analysis revealed a substantial decline in membranal and cortical expression of both GLUT4 and MYH10, where the MCR was decreased 1.7-fold for GLUT4, and 2-fold for MYH10 in the presence of the inhibitor (Fig. 8D-E). Taken together, these results suggest that PKCζ inhibition attenuated cortical MYH10 assembly and GLUT4 translocation to the membrane. Notably, the PKCζ inhibitor also affected the formation of the GLUT4:MYH10 complex, with fewer complexes formed in the inhibited cells compared to the insulin-induced cells (Fig. 8F). Overall, our results demonstrate the importance of MYH10 in adipocyte function and adipogenesis through its involvement in insulin induced GLUT4 translocation regulated by PKCζ. These results provide important insights into the relationship between insulin signaling, GLUT4 translocation, and cytoskeleton rearrangement, where MYH10 play an essential role.

## Discussion

Differentiation of preadipocytes is a complex process that is associated with changes in cell morphology. The early stages of adipogenesis require extensive remodeling and organization of various cytoskeletal components regulated by changes in the ECM^52,57,58^, while in latter parts of differentiation and general metabolism, these components are key in maintaining the physiological function of the cells^5,11^. Nevertheless, our knowledge of such elements and the molecular pathways by which they affect the cells, remains unclear. This study establishes the importance of MYH10 in adipogenesis and adipocyte function, and more specifically, its interaction with GLUT4 and facilitation of translocation of GLUT4 to the cell membrane as part of the insulin signaling pathway.

We initially identified changes in the distribution of MYH10 filaments during the course of adipogenesis. Non-muscle myosins (NMMs) are tightly related to actin, and their interaction is known to affect rudimental cellular processes, including morphogenesis and cytokinesis^59–61^. The cytoskeleton is vital for determining and maintaining the shape and function of adipocytes, and depolymerization and repolymerization of actin affect both early adipogenesis and terminal differentiation^7,58^. Notably, reorganization of other cytoskeletal components has been shown to affect lipogenesis, mitochondrial activity, and glucose uptake in adipocytes ^13–15,62,63^. MYH10 filaments were shown to undergo reorganization in a similar fashion to actin filaments^6^, with a stress-fiber-like appearance in preadipocytes, in contrast to the cortical distribution seen in differentiated cells. These findings further support the notion of reorganization of cytoskeletal components as a potential prerequisite step in adipogenesis. They also highlight the association between myosin and actin in cytoskeletal reorganization and the possible importance as regulators of differentiation.

Subsequently, we established a knockdown model of MYH10 in 3T3-L1 cells to examine how depletion of the protein affects preadipocyte and adipocyte functions. First, we assessed the consequences for migration and adhesion in preadipocytes. MYH10 plays an essential role in cell polarization, motility, and migration through the assembly of actomyosin filaments^31,32^. Previous in-vivo and in-vitro studies have demonstrated the importance of MYH10 in migration. Impaired motility was reported in MYH10 depleted lung carcinoma cell line, glioma cells and mouse embryonic fibroblasts^64,65,66^. In-vivo, MYH10 knockout mice exhibited impaired development of cardiac and brain tissue due to impaired cell adhesion and migration^67–69^. Our observations of a significant impact of MYH10 knockdown on cell migration and spreading are consistent with those of previous studies.

A major finding of this study was that MYH10 has a significant effect on adipogenesis, with knockdown cells displaying no morphological or molecular indications of adipogenesis. The function of MYH10 as a regulator of differentiation can probably be attributed to the interaction with other cytoskeletal components such as actin, that together with ECM signaling and modifications, regulate differentiation. Previous studies have reported the importance of MYH10 in the morphogenesis of a variety of organs through regulation of cell shape and ECM remodeling^70–72^. However, our study is the first to show the importance of MYH10 in adipocytes as prior research focused mainly on MYH9, a paralog of MYH10 and a member of the NMM family^25,26,29,37,38,73–75^. Such studies reported that MYH9 affects the secretion of adiponectin and the translocation of GLUT4 vesicles through an interaction with actin filaments^26,38,75^. The relationship between MYH9 and GLUT4 was studied extensively and showed to affect GLUT4 translocation and glucose uptake in insulin-stimulated adipocytes^25,26,29,37,74,75^. Moreover, Blebbistatin, a myosin inhibitor that affects both isoforms of NMMs, also inhibited glucose uptake in insulin-stimulated adipocytes^25,26^. Although the effect was mainly attributed to MYH9, these observations prompted us to examine the relationship between MYH10 and GLUT4.

The results presented here, demonstrate that, both in-vitro and in-vivo, MYH10 and GLUT4 exist in a functional protein complex localized in the cell membrane. In addition, they also implicate the involvement of MYH10 in the insulin pathway since induction with insulin altered its localization and interaction with GLUT4. Since previous studies reported that, in contrast to MYH9, MYH10 is highly expressed in the cortex of both stimulated and stimulated adipocytes^26^, we used the MCR method to assess the changes in its cortical expression to reveal the upregulation after exposure to insulin. The same method was also used to quantify the MCR as an indicator of membranal expression of GLUT4^49^. The results indicate that MYH10 and GLUT4 are upregulated in response to insulin, both individually and as a complex. We therefore suggest that the insulin dependent GLUT4 translocation to the cell membrane, may be regulated by MYH10. These results may help us to better understand the overlapping and unique roles of different NMMs, specifically in adipocyte function, and the importance of MYH10 in adipogenesis.

The MYH10 knockdown adipogenesis model showed that the WT population in the cultures can induce the differentiation of KD cell, with several differentiated GFP^+^ cells primarily in areas with differentiated WT cells. This finding suggests that the WT population may secrete and transfer factors that induce adipogenesis even in MYH10-KD cells. Because of the observed interaction between GLUT4 with MYH10, we hypothesized that co-culturing of MYH10-KD with GLUT4^+^ cells could have an effect on the adipogenic potential of the MYH10-KD cells. In this context, adipocytes are known to be able to sense their niche and interact with neighboring cells either directly or indirectly^76,77^. Notably, we were able to restore some adipogenic capacity to MYH10 depleted cells by uptake of GLUT4 particles from neighboring GLUT4^+^ cells in co-culture (Fig. 7C,E). The transport of extracellular vesicles containing GLUT4 from other cells, has been reported previously^78^, and this method of cellular communication is of great interest because of its potential in the regulation of adipocyte differentiation and function.

The regulation of NMMs differs from that of cardiac and muscle myosins and involves the phosphorylation of regulatory light chains and the tails of the heavy chains themselves^30,32^. Phosphorylation of the non-muscle myosin tails can promote the reorganization and localization of actomyosin filaments^79–84^. We predicted that some regulators of MYH10 may also be downstream effectors of the insulin pathway and may govern MYH10 activity in that regard. We were able to identify PKCζ as a regulator of MYH10 function in induced adipocytes. Our investigations into potential regulators of MYH10 activity indicated a possible role for PKCζ in regulating MYH10 activity via the insulin pathway, and indeed, its inhibition impeded MYH10 and GLUT4 activity. PKCζ is an atypical protein kinase C protein that was previously shown to be highly related to the insulin pathway. It is phosphorylated by phosphatidylinositol (PI) 3-kinase and, in turn, can phosphorylate various downstream effectors that regulate GLUT4 translocation^39,40^. PKCζ also regulates the required cytoskeletal reorganization that accompanies insulin signaling and can affect the polymerization of actin that is crucial for GLUT4 shuttling, placing it at the intersection of insulin signaling and cytoskeleton activity^85–87^. The different non-muscle myosins have several overlapping and unique regulators both through their regulatory light chains and heavy chains^30,32^. Interestingly, MYH10 is the only non-muscle myosin regulated by PKCζ, which highlights the differences in function and regulation of the different myosin isoforms^79,88,89^. The reported effects of PKCζ on the cytoskeletal association, mechanoresponsiveness, and cortical localization of MYH10 are in good agreement with our finding that it regulates the localization of MYH10 in response to insulin and that inhibition of PKCζ inhibited MYH10 and GLUT4 activity^79,90^. Our observations suggest that PKCζ is phosphorylated as part of the insulin pathway, and in turn, can trigger MYH10 cortical activity that facilitates GLUT4 translocation to the membrane because of their interaction (Fig. 8G). The suggested pathway further highlights the strong relationship between cytoskeleton activity and cellular functioning.

In conclusion, we have identified MYH10 as a novel effector of adipogenesis and adipocyte function. Our results demonstrate that MYH10 regulates the translocation of GLUT4 through insulin-induced PKCζ activation, and that MYH10-KD inhibits adipogenesis. These observations further support the importance of cytoskeleton proteins in adipocyte function and differentiation. Future in-vivo studies that incorporate pathophysiological conditions and their effect on MYH10 function in adipocytes will undoubtedly prove informative.

## Acknowledgments

Mike Egozi participated in the project as part of the jInternship program. We acknowledge Ann Avron for the editorial assistance.

